# Microbial dispersion in the human gut through the lens of fecal transplant

**DOI:** 10.1101/2023.08.29.555267

**Authors:** Yadid M. Algavi, Elhanan Borenstein

## Abstract

Microorganisms frequently migrate from one ecosystem to another, influencing and shaping their new environment. Yet, despite the potential importance of this process in modulating the environment and the microbial ecosystem, our understanding of the fundamental forces that govern microbial migration and dispersion is still lacking. Moreover, while theoretical studies and in-vitro experimental work have highlighted the contribution of biotic interactions to the assembly of the community, identifying such interactions in vivo, specifically in communities as complex as the human gut, remains challenging. To this end, we developed a new, robust, and compositionally invariant approach, and leveraged data from well-characterized translocation experiments, namely, clinical fecal microbiota transplant (FMT) trials, to rigorously pinpoint dependencies between taxa during the colonization of human gastrointestinal habitat. Our analysis identified numerous pairwise dependencies between co-colonizing microbes during migration between gastrointestinal environments. We further demonstrated that identified dependencies agree with previously reported findings from in-vitro experiments and population-wide distribution patterns. Finally, we characterized the web of metabolic dependencies between these taxa and explored the functional properties that may promote better dispersion. Combined, our findings provide insights into the principles and determinants of community dynamics following ecological translocation, informing potential opportunities for precise community design.

## Introduction

Dispersion, the distribution of taxa from one habitat to another, is a crucial process that allows microbes to colonize new ecosystems, interact with proximate co-existing communities, and transfer genetic material and metabolic byproducts^1^. This fundamental community assembly process shapes the natural environment, from aquatic and terrestrial ecosystems^2–4^ to host-associated communities^5^. Within the human microbiome, dispersion is critical for maintaining biodiversity on both spatial (e.g., migration from proximate biogeographic niches through the digestive tract^6^) and temporal (e.g., recurrent waves of invading microbes in the infant gut^7,8^) scales. There is even some evidence that non-communicable diseases, such as cancer and cardiovascular disorders, may be transmitted by microbial components^9^, highlighting the importance of understanding the role of migration in community assembly.

From the perspective of the dispersing taxa, to establish a viable population, it has to adapt to the unique conditions in the new habitat, including nutrient availability in this new environment, co-existing competing taxa, and, in host-associated environments, elements of the host immune system^10,11^. Moreover, as bacteria rarely exist in isolation, migration could be portrayed as the mixing of two communities^12^. Indeed, the significance of community-level interactions for the assembly of mixed communities has been highlighted by both mathematical models^13^ and experimental studies^14–16^. In the context of the human microbiome, microbial dependencies during dispersion can be studied via naturally occurring host-host interactions. For example, some work has studied the migration of bacteria across human hosts in a shared household^17^, during contact sport^18^, or via intimate kissing^19^. Notably, however, the scarcity of such data, and the stochastic nature of such interactions, pose various experimental challenges and limit our ability to rigorously study microbial dependencies during translocation. More recently, a new opportunity for examining the dispersion of whole gut microbial populations between hosts has emerged, owing to the increased prevalence of fecal microbiome transplants (FMTs)^20–22^. Indeed, FMT-based data have been utilized to characterize taxa replacement and engraftment efficiency and to explore the medical consequences of such treatments. Yet, while FMT clinical trials facilitate a more controlled and well-defined study setup, the compositional nature of metagenomic data and the statistical challenges it induces, generally limit the ability to infer microbial interactions beyond presence-absence patterns^23^. As a result, many fundamental ecological aspects concerning dispersion in the human gut remained poorly-characterized, such as which taxa are more easily dispersed, what is the role of the potential dependencies between co-migrated taxa, and what causes certain taxa to become dominant in the new host.

In this study, we wish to address these questions by developing a statistically robust measure that we term *Relative Dispersion Ratio (RDR)*, which aims to reveal pairwise interactions between co-colonizing taxa. Using this measure, we analyzed data from a meta-cohort of 8 FMT studies, identifying, cataloging, and characterizing co-colonizing taxa dependencies. We further compared identified interactions with both in-vitro pairwise experiments and population-wide distribution patterns, demonstrating that these interactions play an important role in microbiome assembly. Lastly, we pinpoint the metabolic and functional drivers that may lead to superior dispersion. Overall, our findings provide valuable insights into the mechanisms driving microbial migration and the role of biotic interactions in shaping and transforming the environment.

## Results

### The Relative Dispersion Ratio (RDR) – A robust measure of successful dispersion

In migration scenarios, colonization patterns can provide intriguing insights into microbial assembly dynamics and into the success of colonizing taxa in the new environment and its determinants. To characterize such colonization patterns, we therefore utilized publicly available FMT data. In these data, samples are arranged in triads, with each triad including one donor sample before FMT, and a pair of recipient samples, one before and one after FMT (Figure 1A). In this work, we refer to each such triad as an *FMT experiment*. In this setting, *colonizers* can be defined as taxa that appeared in both the donor and recipient post-FMT samples but were absent from the recipient pre-FMT sample^24^. Such microbes migrate during FMT from their source environment (the donor’s gut), and get established in a new environment (the recipient’s gut), in which they were not previously present. In this new host environment, they may face markedly different conditions, community composition, and niche availability, which in turn may determine how abundant they may ultimately become in this host. Our analysis focuses on these taxa, as a vehicle to reveal how microbes adapt and disperse when migrating to a new ecological niche.

**Figure 1.**
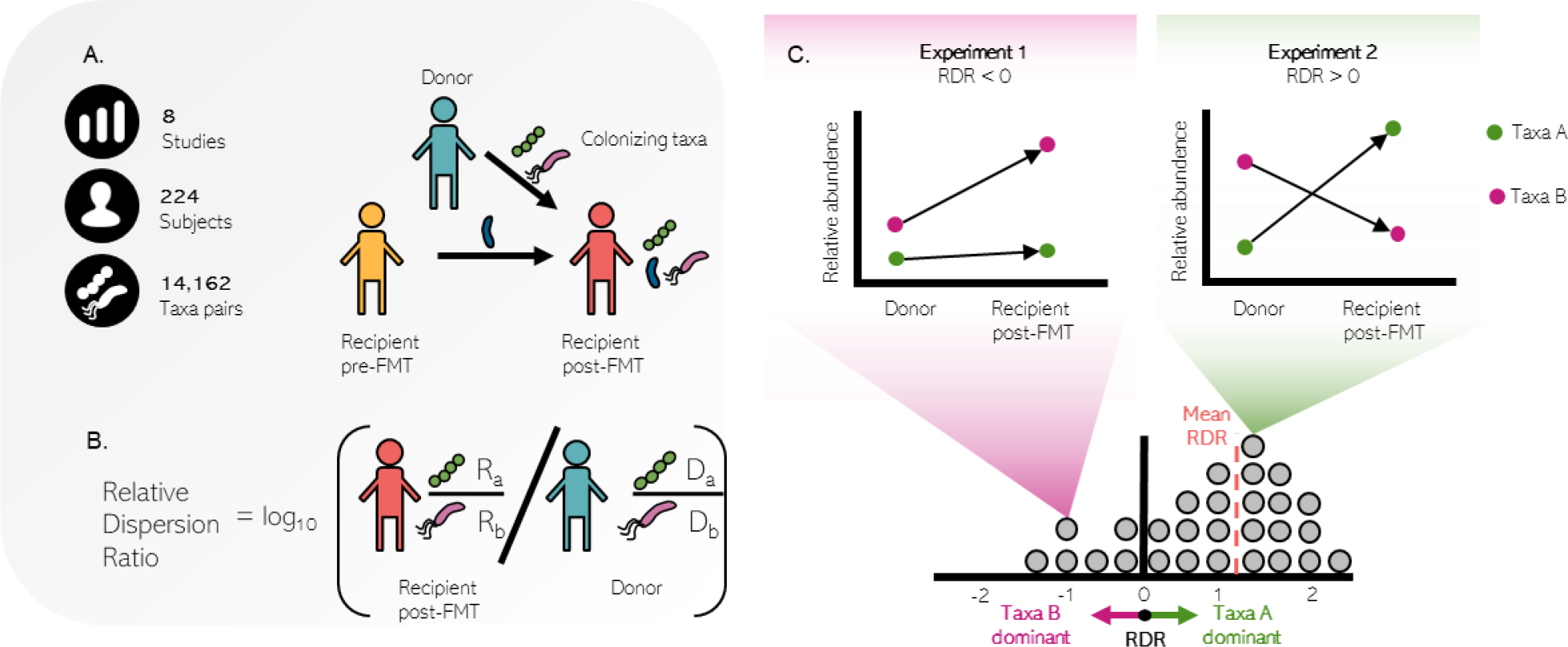
Community structure and dynamics following fecal microbiome transplant treatment. **A**–Data structure and scale of our FMT dataset collection. Each FMT procedure can be viewed as a mixing of two microbial populations, the donor and recipient pre-FMT, which eventually results in a single recipient post-FMT community. **B**–RDR is defined as the fold change of the ratio between the relative abundances of two co-colonizing taxa in the new host (recipient post-FMT) and the former host (the donor). **C**–Illustration of possible FMT dynamics as captured by the RDR framework. RDR > 0 indicates relatively better dispersion of taxa A compared to taxa B during migration from the donor to the recipient environments (right top panel), and vice versa for RDR < 0 (left top panel). For each pair of taxa, we can calculate RDR across all FMT experiments in which they appear as co-colonizers, resulting in a distribution of RDR scores (bottom panel). This distribution can be used to calculate the mean RDR (red dashed line), as well as various statistics.

Notably, however, due to the compositional nature of metagenomics data, which provide information about the relative abundance of each taxon in the community rather than its absolute abundance, the fold change between a taxon’s abundance in the source environment and its abundance in the new environment cannot rigorously attest to its success in this new environment^23^. We therefore introduce below a more robust metric, termed *Relative Dispersion Ratio (RDR)*, which is invariant to compositionality, thus allowing us to uncover complex interactions between colonizing taxa and their ecology more confidently. Specifically, the RDR metric aims to quantify the difference in dispersion between two microbial taxa, and is defined as the ratio of abundances of each of two co-colonizing taxa between the two environments (Figure 1B). Formally, assuming *a* and *b* represent two co-colonizing taxa in some FMT experiment, with *D*_*a*_ and *D*_*b*_ denoting their relative abundances, respectively, in the donor, and *R*_*a*_ and *R*_*b*_ denoting their relative abundances in the post-FMT recipient, then we define *RDR* as *log*_*10*_*[(R*_*a*_*/R*_*b*_*)/(D*_*a*_*/D*_*b*_)]. Focusing on the RDR of pairs of co-colonizing taxa thus allows us to quantify which bacteria are more dispersive *compared to the other* in the new environment of the recipient host. Put differently, RDR>0 can be interpreted as indicating a more successful dispersion of the first taxa in compression to the second, while RDR<0 indicates the opposite (i.e. the second taxa dispersed better than the first; Figure 1C, top panels). Throughout the text, we will name the better-dispersed taxon in a pair of co-colonizing taxa the *dominant partner*, and the worse-dispersed taxa the *minor partner*. Note that when RDR>0, for example, it is still possible that both taxa increased in absolute abundance, that both decreased in absolute abundance, or even that both decreased in relative abundance post-FMT, and that indeed, RDR can only attest to the *relative* success of the two taxa. Importantly, however, since this metric is based on ratios between relative abundances, RDR is not affected by the compositionality of microbiome data and does not suffer from spurious statistical properties as standard analyses of relative abundance data. Finally, when a pair of taxa appear as co-colonizers in multiple FMT experiments, their RDR can be calculated in each experiment independently, resulting in a distribution of observed RDR scores for these two taxa, as well as their *mean RDR* (Figure 1C, bottom panel).

### RDR identifies dispersion differences in FMT settings

We next sought to apply this metric to a large set of FMT samples, aiming to characterize dispersion patterns. To this end, we first constructed a dataset collection obtained from 8 clinical FMT studies. This meta-cohort covers multiple underlying diseases, including three studies of patients with *Clostridium difficile* infections^25–27^, three with inflammatory gut disorders^28–30^, one with autism spectrum disorder patients^31^, and one with melanoma patients^32^. Combined, these datasets thus represent the diversity of the patient populations commonly studied in FMT research. In total, these datasets include 624 16s rRNA metagenomic samples from 124 patients and 100 healthy donors (Supplementary Table 1). We treated each triad of related samples (donor pre-FMT, recipient pre-FMT, and recipient post-FMT) as an FMT experiment, resulting in a total of 356 such experiments. Although all studies sequenced the V4 or V3-V4 regions, we re-analyzed the raw sequences using QIIME2^33^, to reduce variability in downstream analysis. We further profiled each metagenomic sample at the genus level using the GTDB hierarchy^34^ and identified 201 unique genus-level taxa (see Methods).

As noted above, to explore how changing environment impacts taxa prevalence, we primarily focused on colonizing taxa. We found that on average each FMT experiment introduces 31±20 new taxa into the recipient ecosystem (Supplementary Figure 1A), suggesting that colonizing taxa do not disperse in isolation or independently of other taxa. Furthermore, these colonizing taxa form a substantial portion (32±22%) of the post-FMT overall community abundance (Supplementary Figure 1B). Interestingly, examining the compositions of donor and recipient samples and cataloging all colonizer taxa and their abundances, we further found that 90% of the colonizing taxa undergo on average at least two-fold variation in their relative abundance following FMT (either two-fold increase or two-fold decrease; Supplementary Figure 1C), although, as described above, this does not necessarily mean that they increase in absolute abundance.

We finally used the data above to calculate RDR for 14,162 taxa pairs that appear as co-colonizing taxa in at least one FMT experiment. Importantly, however, many of these pairs appear, as co-colonizing taxa, in multiple FMT studies and in multiple FMT experiments (mean 2.2 studies and 7.2 experiments). To focus on pairs of taxa for which the consistency of co-colonization patterns can be quantified confidently, we discarded from our analysis pairs of taxa that were not highly prevalent, considering only the 3,160 taxa pairs that appeared as co-colonizing taxa in at least 2 studies and 10 experiments (which we term *prevalent pairs*). We then performed a one-sample Wilcoxon signed-rank test to identify pairs for which the calculate RDR across the various experiments in which they appear is significantly different from 0 (with FDR-corrected p-value<0.05), indicating that one member of the pair dispersed significantly better than the other. Our analysis identified 942 such pairs, which we will term *differentially dispersing pairs* throughout the text. For each such pair, we assess the magnitude of differential dispersion by calculating the mean RDR across all experiments in which the pair co-colonized, and the identity of the dominant partner by the sign of the mean RDR (i.e., RDR>0 indicates that the first species, *a*, is the dominant taxon, and RDR<0 indicates that the second species, *b*, is the dominant partner; see Figure 1B). In total, these pairs represent eight phyla and over 100 genera. Figure 2A-D presents the observed colonization patterns for ten representative differently dispersing pairs, including their RDR score in each experiment and mean RDR (Figure 2A), statistical significance (Figure 2B), and the number of FMT experiments and studies in which they were found (Figure 2C,D).

**Figure 2.**
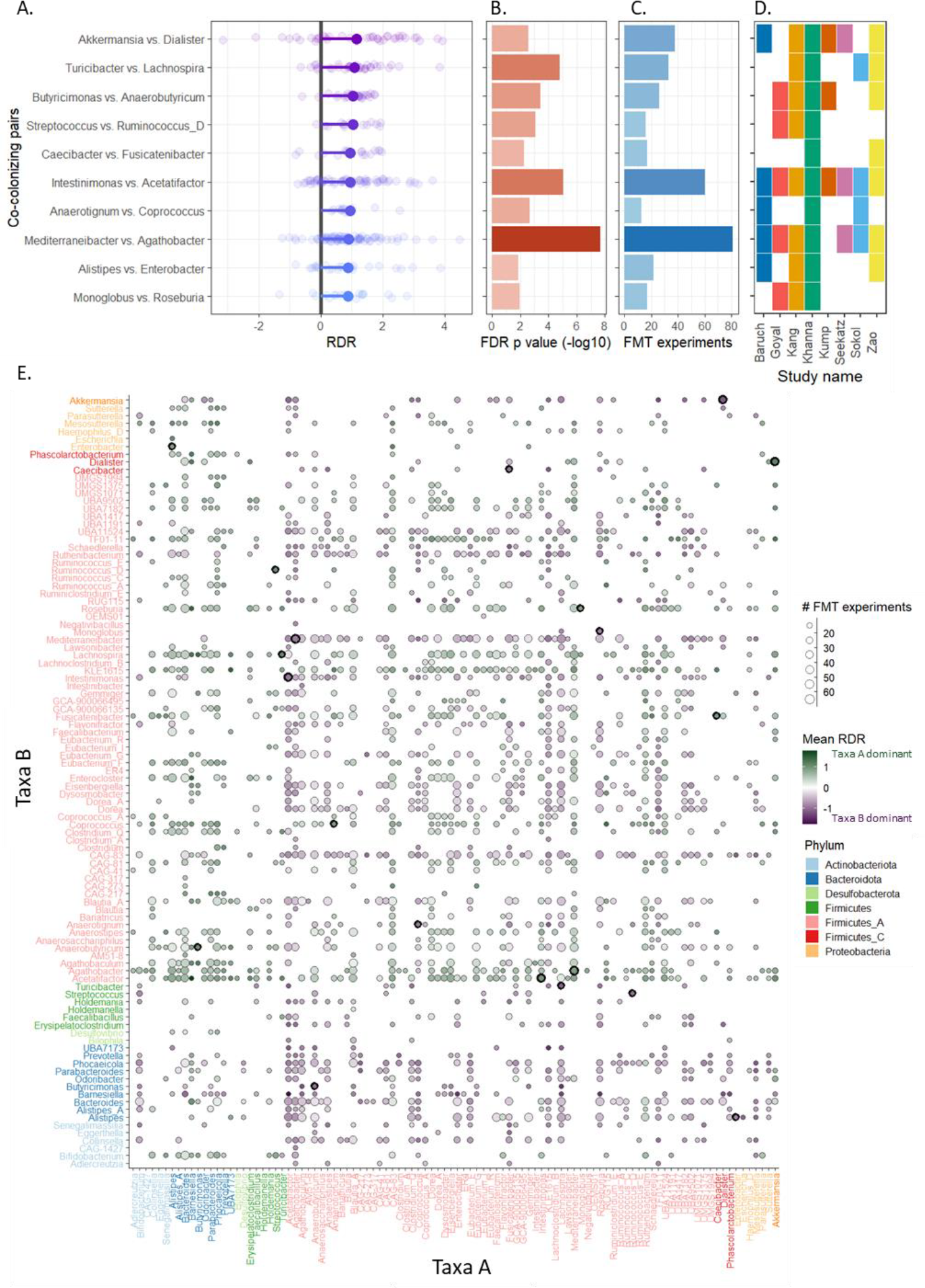
Relative Dispersion Ratio (RDR) scores across differentially dispersing pairs. **A**–Each dot represents the calculated RDR from a single FMT experiment, with the large dot representing the mean RDR. For convenience, we ordered each pair such that the dominant partner is the first taxa and the minor partner is the second species (and hence, all mean RDR values are positive). **B**–FDR-corrected p-values of one-sample Wilcoxon signed-rank test. **C**–Number of FMT experiments in which the pair co-colonized. **D**–Identity and number of studies in which the pair appeared. **E**– The complete mean RDR table between all differentially dispersing pairs. The color of each dot indicates the identity of the dominant partner (Green.taxa A on the X axis, Purple.taxa B on the Y axis). Dot size represents the number of FMT experiments in which the pair of taxa co-colonized. The color of the labels on the X and Y axis indicates the phylum. Black outer circles denote the taxa pairs included in panel A.

To confirm and demonstrate the generalizability of our findings within the context of FMT studies, we further used a leave-one-study-out cross-validation approach. Specifically, we removed one study from the meta-cohort each time, and for each of the 942 differentially dispersing pairs described above, compared the mean RDR of the pair in this study alone, to the mean RDR calculated for the seven remaining studies. We found that the sign of the mean RDR (and hence the identity of the dominant partner) is the same in more than 95% of these leave-one-out calculations, indicating that our RDR analysis is not study-specific.

The complete table of mean RDR scores for all differentially dispersing pairs is available in Fig 2E. Inspecting the identity of the dominant and the minor partners in these pairs, we found that members of the phyla *Bacteroidota* and *Actinobacteriota* frequently tend to be the dominant partner, followed by *Firmicutes* and *Proteobacteria* that are more often the minor partner (Supplementary Figure 2, Wilcoxon test). This is in line with previously reported phylum-dependent colonization rates during FMT^20–22^. Among genera with high dominance rates, are prevalent members of the human gut microbiome such as *Bacteroides* and *Prevotela*, as well as multiple members of the family *Lachnospiraceae*, including *Dorea* and *Blautia*. Indeed, these taxa have been well-documented for their ubiquity in various environments^35–38^.

### Comparing RDR scores to results from in-vitro experiments and population-wide distribution patterns

To confirm the dispersion differences inferred by the RDR methodology above, we compared them to findings from in-vitro experiments in which distribution differences between human microbiota members could be measured explicitly. To this end, we obtained data from a study of 36 pairwise in-vitro competition experiments, in which two gut-dwelling strains were incubated in a co-culture for 72 hours while their abundances were monitored^39^. While these experimental settings are clearly markedly simpler in comparison to FMT (in-vitro vs in-vivo, co-culture vs community, etc.), we still expect dominant partners to generally gain higher abundance in the co-culture at the end of the experiment. Indeed, we found a high degree of agreement between the results of our RDR framework and the in-vitro experiments (Figure 3A). Specifically, considering the 9 differentially dispersing pairs included in this in-vitro study, we found that 8 (89%) agreed on which of the two taxa was dominant. Furthermore, examining all taxa pairs included in this in-vitro assay (i.e., including those for which our analysis did not reach significance and hence were not identified as differentially dispersing pairs), we found that in 25 out of 36 comparisons, the mean RDR and in-vitro experiments again agreed on the dominant taxa (69% of interactions, Cohen’s kappa, κ= 0.39, p= 0.018).

**Figure 3.**
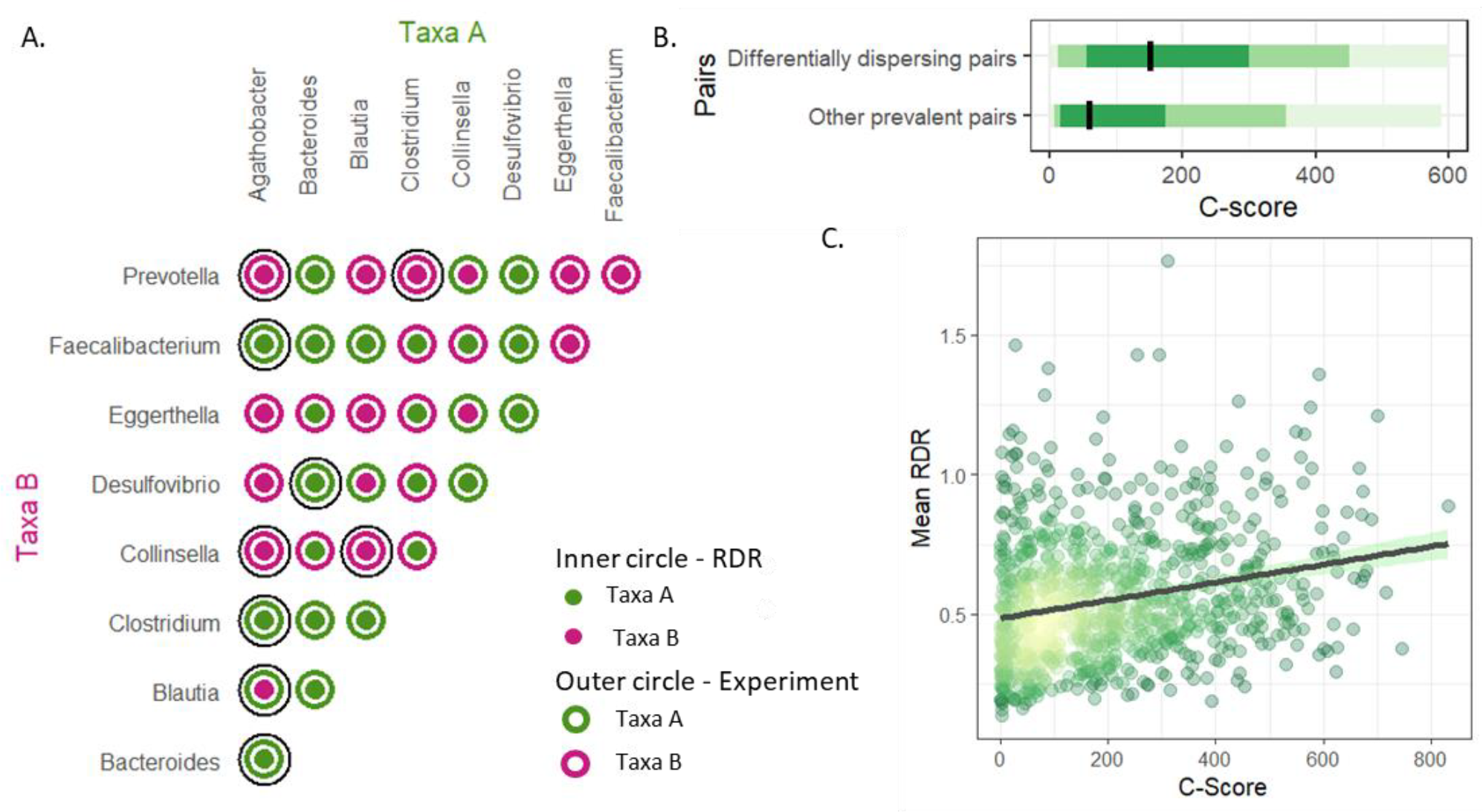
Comparison of RDR to observed patterns in in-vitro experiments and population-wide distributions. **A**–Comparison between the identity of the dominant taxon as inferred from the RDR framework and from in-vitro competition experiments. Each dot represents a single in-vitro competition experiment between a pair of microbes. The color of the outer and inner circles indicates the identity of the dominant taxa in the in vivo experiment and RDR pairs, respectively (green indicates that the dominant taxon is the one listed on the x-axis while purple indicates that the dominant taxon is listed on the y-axis). Matching inner and outer colors indicates agreement between the RDR and experiment. The outer-most black circles highlight the 9 differentially dispersing pairs **B**–The difference in C-score across the American gut project (AGP) cohort between significant RDR pairs and other prevalent pairs (two-tailed Wilcoxon test, p=3.32*10^-27^). Different shades of green indicate 50%, 80% and 95% confidence intervals. **C**–Correlation between C-scores from the AGP cohort and mean RDR scores for all differentially dispersing pairs (Spearman correlation, ρ = 0.25, p= 1.83*10^-5^).

To further assess the relevance of our RDR framework for characterizing community assembly, we next examined taxa distribution patterns across human populations. While this setting likely captures additional dimensions of environmental selection beyond differences in dispersion capabilities (such as interactions with other gut-dwelling microbes, host nutrition, and health conditions), it could serve as a proxy for the long-term outcome of such differential dispersion dynamics. To this end, we turned to the American Gut Project (AGP), a cohort containing metagenomic samples from nearly 10,000 participants^40^. To minimize bias due to differences in metagenomic processing pipelines, we re-analyzed the raw sequencing data and annotated it according to the GTDB taxonomy (Methods). We then calculated the checkerboard score (C-score), which is a measure of pairwise taxa distribution across different habitats, characterizing competition among taxa and resulting taxa segregation^41^. We found that differentially dispersing pairs exhibited significantly higher C-scores in comparison to the C-scores of all other prevalent pairs, potentially suggesting the existence of exclusion dynamics between these taxa during community assembly^42,43^, which is in line with their differential dispersion success identified via our RDR metric (Figure 3B, Wilcoxon test p=3.32*10^-27^). Interestingly, we further found that the C-score of these pairs is significantly correlated with their mean RDR (p =1.83*10^-15^; spearman correlation test), suggesting that differences in dispersion capabilities are associated with population-wide taxa segregations patterns (Figure 3C). Taken together, these analyses suggest that the relative dispersion differences inferred via the RDR framework are well mirrored in both in-vitro experiments and population-wide distribution patterns.

### Establishing RDR dispersion patterns based on mechanistic metabolic model-based methods

We next set out to identify plausible mechanistic determinants of the observed dispersion differences between the two members of each differentially dispersing pair. We first resorted to a metabolic network-based analysis, using specifically a previously introduced framework, termed reverse-ecology^44^,which was shown to successfully predict competitive and cooperative interactions between microbial species, as well as ecological design strategies^45,46^.

Briefly, for this analysis, we first constructed genus-level metabolic networks, using genomic representation from PICRUST2^47^ (Methods). We then applied the seed-set detection algorithm^45,48^ to infer the organism’s nutritional profile and used the fraction of seed metabolites from the total number of metabolites in the network as a proxy for the nutritional flexibility of this genus and its ability to flourish across multiple biochemical habitats as previously suggested^49^. Comparing this measure to our calculated mean RDR, we found that dominant partners had on average a higher nutritional flexibility (Figure 4A, two-tailed Wilcoxon test, p=1.06*10^-5^), which can be beneficial for adapting to different metabolite availability in a new environment. We also used the identified seed sets to compute the metabolic competition index between pairs (following Levy at al^45,49^). This index was shown to provide a proxy for niche overlap and estimates the potential level of competition one taxon may experience in the presence of the other. Compellingly, we found that the dominant partner experienced significantly less competition in the presence of the minor partner than the minor partner did in the presence of the dominant partner (Figure 4B). Together, the increased metabolic flexibility and the lower competition may thus indicate an enhanced metabolic capacity of the dominant partner compared to its minor partner.

**Figure 4.**
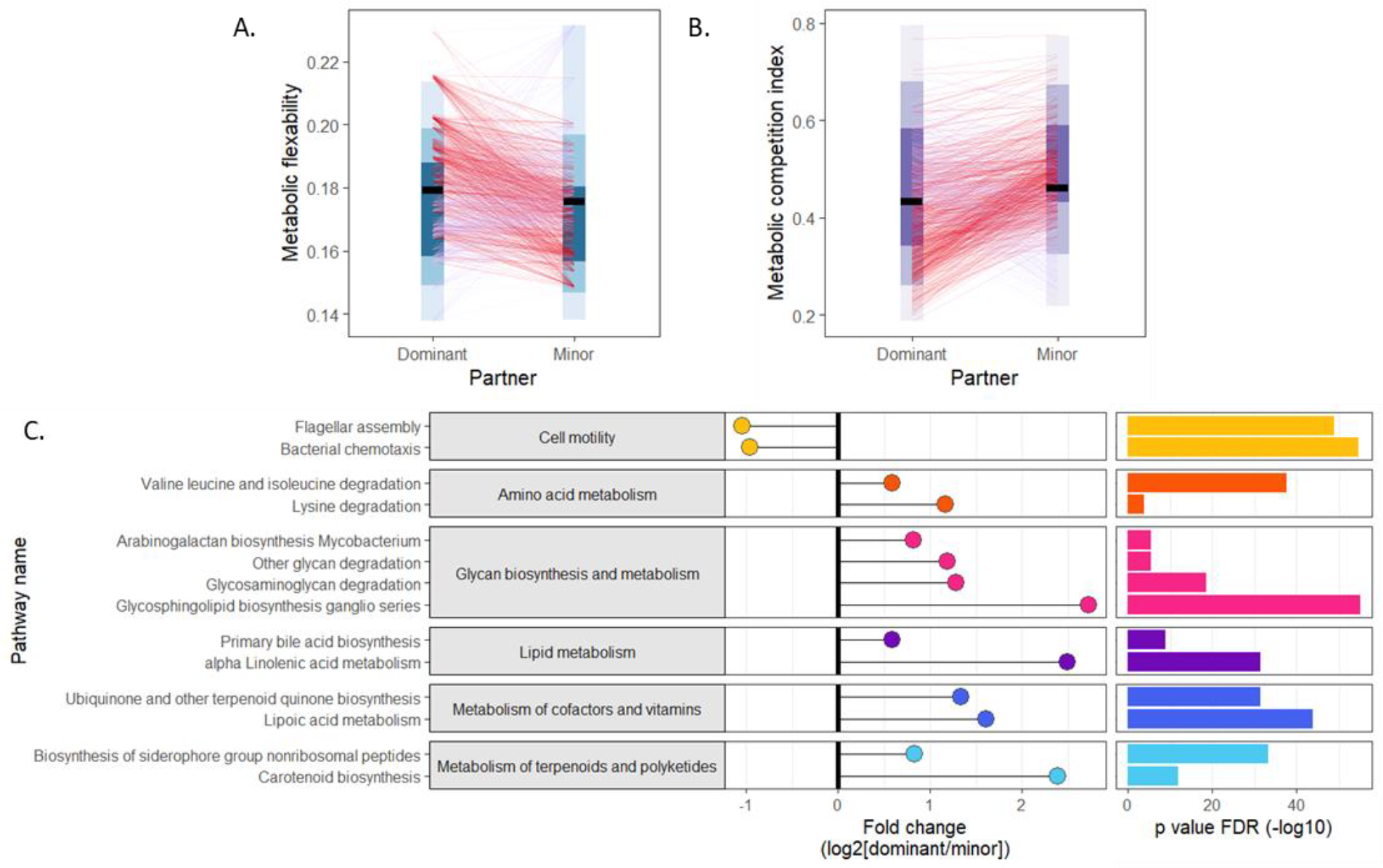
Comparison of the metabolic capacity of the dominant vs. minor partner in differentially dispersing pairs. **A**–Pairwise comparison of metabolic flexibility (calculated based on the seed-set framework; Methods) between the dominant and minor partner (paired Wilcoxon signed-rank test, p=1.06*10^-5^). Different blue shades indicate 50%, 80% and 95% confidence intervals. **B**–Pairwise comparison between the metabolic competition index between the dominant and minor partner (paired Wilcoxon signed-rank test, p=6.59*10^-10^). Different purple shades indicate 50%, 80% and 95% confidence intervals. **C**–Significant differences in bio-synthetic capacities between dominant and minor partners. Pathways are organized according to pathway categories. Positive values indicate enrichment among dominant taxa while negative values indicate depletion.

Finally, we set out to examine whether differences in dispersion success as measured by the RDR framework are also associated with differences in biosynthetic and functional capacities between the two members of each differentially dispersing pair. To this end, we compared the number of KEGG KOs in each pathway between the members of each such pair to identify pathways with significantly more or less genes in the dominant vs. the minor partner (Methods). Our analysis identified 14 such KEGG metabolic pathways, spanning multiple functional categories (Figure 4C). Specifically, this analysis suggested that highly dispersive taxa tend to encode fewer cell motility genes. Indeed, cell motility associated genes are known to have high energetic costs and cause reduced growth rates^50^. In addition, these motility genes have also been associated with lower engraftment rates during FMT^20^. In addition, we found that dominant taxa tend to have a variety of enriched biosynthetic capabilities, including the metabolism of essential amino acids such as valine, lysine, and isoleucine, which have been identified as important factors in the assembly of microbial communities^51^. Dominant taxa were further associated with enrichment of pathways for glycans breakdown, which serve as energy sources for gut microbes. Indeed, the ability to utilize a variety of glycans and dietary fibers has been shown to expand the microbe’s ecological niche in a new habitat^52^. Moreover, we observed enrichment in pathways for synthesizing cofactors involved in energy production, such as lipoic acid, used in the TCA cycle, and ubiquinones, redox-active compounds during ATP production^53^. Similarly, bile acid transformation pathways, including those for primary bile acid and alpha-linolenic acid, were also enriched among dominant taxa, in agreement with the known impact of bile acid repertoire modulation on gut microbiome and host health^54^. Finally, there was an increase in the metabolism of non-ribosomal peptides, which have crucial roles as siderophores and virulence factors^55^. Overall, this analysis has identified several potential functional factors that may contribute to successful dispersion.

### Partners in differentially dispersing pairs are highly informative for predicting post-FMT abundances

Given the observed consistency of relative dispersion rates in differentially dispersing pairs, and more importantly, the potential metabolic determinants underlying the observed dependencies between co-colonizing taxa, we next examined whether such pairs can also be linked to our ability to predict community structure after FMT. To this end, we used random forest regressor models to predict the relative abundance of every taxon post-FMT based on pre-FMT community compositions in the donor and recipient (Methods). We evaluated the performances of these models using Spearman correlation between the predicted and observed abundance, and defined taxa that exhibit FDR corrected p-value < 0.05 and Spearman ρ > 0.3 as *well-predicted*. Notably, out of the 100 taxa analyzed, 52 were well-predicted with a mean R^2^ of 0.21 (Figure 5A). Focusing on these well-predicted taxa, we quantifies the contribution of each pre-FMT taxa in the donor and in the recipient to the model using Shapley additive explanations (SHAP) values^56^. For subsequent analysis, in evaluating each predictive model, we partitioned the pre-FMT taxa into 3 categories: “Self” – the taxa whose abundance post-FMT the model aims to predict, “Partners” – partners of the predicted taxa in differentially dispersing pairs, and “Non-partners” – taxa that are not partner of the predicted taxa in a differentially dispersing pair. Comparing the SHAP contribution of these 3 taxa categories across the 52 models, we first found, not surprisingly, that the abundance of the predicted taxa (“self”) in the pre-FMT samples has the highest contribution to predicting the taxon’s post-FMT abundance (Wilcoxon test, Figure 5B). However, we further found that partners have significantly higher contribution to predicting post-FMT abundance compared to non-partners (Figure 5B, p = 7.34*10^-9^). Importantly, this trend holds when considering separately pre-FMT abundances in the donor and in the recipient (Wilcoxon test, Figure 5C).

**Figure 5.**
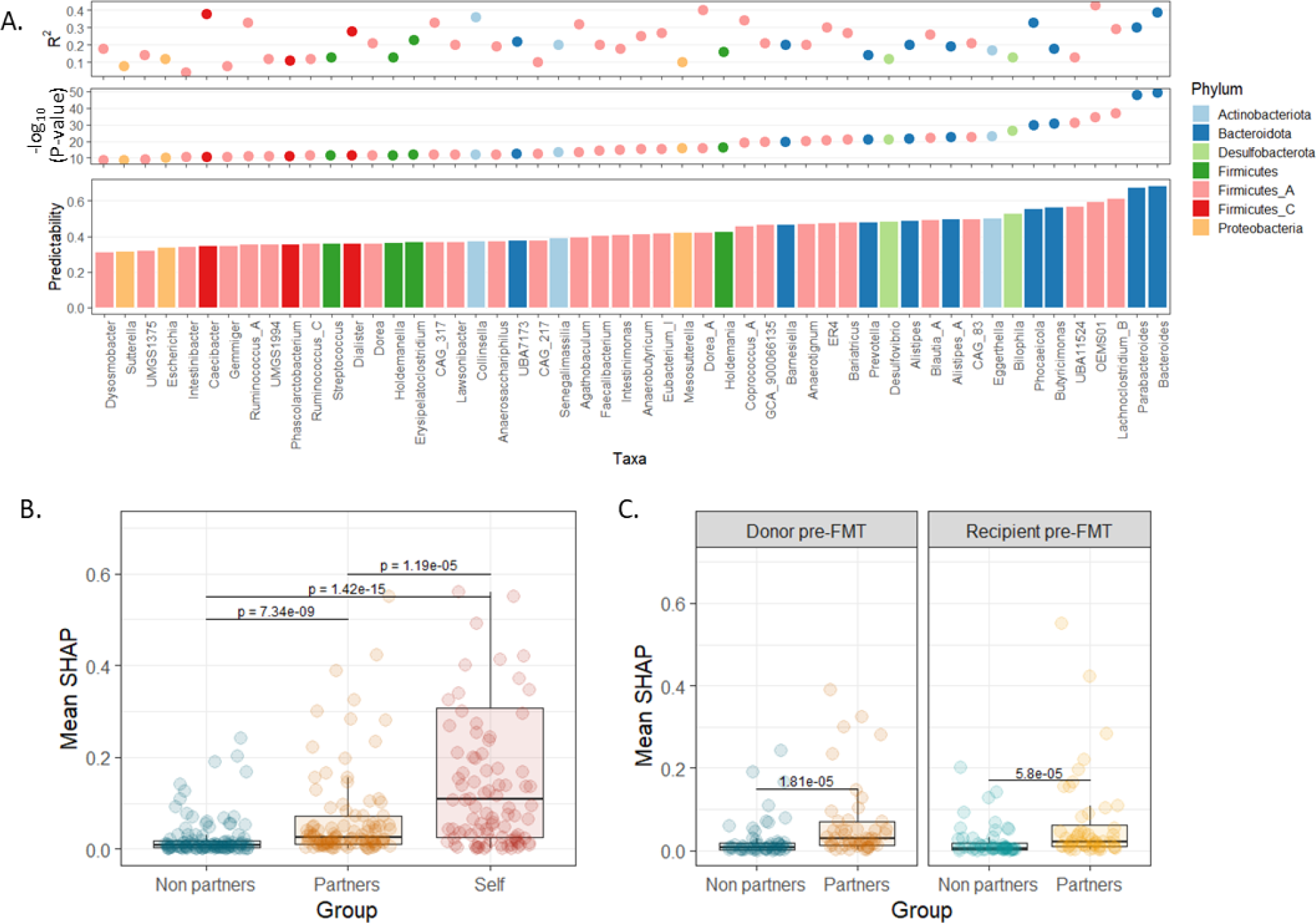
Information gained by co-colonizing taxa for predicting post-FMT community composition. **A**–Machine learning models predict post-FMT abundance. Bottom panel.predictability (measured by Spearman correlation between observed and predicted abundances); Middle panel.FDR corrected p values (in -log10 scale); Top panel.The model coefficient of determination (R^2^). **B**–Comparison between SHAP values of 3 different taxa categories: Self (the pre-FMT abundance of the predicted taxa), Partners (partners of the predicted taxa in differentially dispersing pairs), and Non-partners. C.Comparison between SHAP values (as in Panel B), with separation between donor and recipient profiles.

## Discussion

Understanding the mechanisms underlying the dispersion of microbial communities is an essential, yet poorly understood ecological domain. Previous studies have stated that inter-microbial interactions between invading taxa play a crucial role in modulating the community in various habitats, such as soil, marine environments, and plant roots^2–4^. However, studies aiming to characterize such interactions in human-associated communities are still scarce, owing to various statistical and experimental challenges. These difficulties, for example, have promoted multiple previous studies to analyze dispersion across human gut environments based on presence/absence data only^20–22^ (*i*.*e*., whether the microbe has been detected following FMT or not), ignoring the wealth of information captured in microbiome compositional profiles.

Here, we address some of these challenges and further quantify the outcomes and determinants of microbial dispersion across human gut microbiomes by introducing a new computational framework for analyzing metagenomic data from multiple clinical FMT studies. Our framework aims to uncover consistent differences in dispersion rates, ultimately presenting a web of potential dependencies between migrating gut-dwelling microbes. Importantly, we further provided evidence that identified dispersion differences mirror co-culture experiments and population-wide co-occurrence patterns, and used metabolic modeling and functional enrichment analysis to pinpoint functional and metabolic factors associated with superior dispersion (that were in agreement with reported findings from germ-free mice experiments^57^).

The set of identified co-colonizing taxa pairs that exhibit statistically significant difference in dispersion can be used to better understand potential microbial dynamics during migration, including, for example, underlying competitive or metabolic interactions. However, while our study represents a step forward in understanding the role of biotic and metabolic interactions in dispersion dynamics, it is important to note that these processes are extremely complex and that multiple other environmental factors, such as nutrient avilability^52,58^, host genetic background^59^, the activity of the immune system ^60^, and biogeographic patterns^61^, are all likely playing a role in shaping the composition of the community after fecal transplant. Hopefully, the increase in the prevalence of multi-omics studies of the gut microbiome and of FMT-based data will allow us to further map these complex processes and the contribution of each of these factors to dispersion dynamics.

Notably, from a border perspective, our research aims to tackle a challenging task, inferring how the functional capabilities of individual microbes collectively impact a given community assembly^62,63^. Previous studies of this topic have proposed bottom-up assembly rules, demonstrating that the interaction between individual taxa can explain, at least partly, the observed patterns in community structure. For example, metabolic network analysis^45^ and consumer resource models^45,64^ have proposed that competitive forces are the main drivers of the resulting taxonomic profile. Other works^65,66^, in contrast, have claimed that cooperative cross-feeding interactions are frequent among microbiome members, especially in resource-limiting niches. In the context of microbial migration, specifically, Diaz-Colunga *et al*.^67^ have described the importance of cohesive inter-microbial interactions between members of soil communities, and highlighted the competitive forces between taxa as driving dispersion differences. Our framework, and other analyses of FMT dynamics are similarly essential for uncovering the forces underlying community assembly in the gut, with important and timely implications for microbiome-based therapy.

More generally, we believe that the framework introduced in this work has the potential to be utilized in multiple other settings, where metagenomic data on the origin and destination of microbial migration are available, and could serve as a rigorous method for comprehensively and systematically analyzing microbial dispersion. Looking ahead, our findings, in conjunction with those of other studies, offer exciting opportunities for designing tailored microbial communities to meet the challenges in both environmental and clinical contexts.

## Methods

### Data acquisition and preprocessing

We acquired 16S rRNA amplicon sequencing data from published FMT studies described in Supplementary Table 1. For consistency, we processed and similarly analyzed each dataset. Specifically, we obtained raw fastq files from public repositories (NCBI Sequence Read Archive or European Nucleotide Archive) and processed these data using Qiime2 version 2019-127^68^. We demultiplexed the data using the Qiime2 demux plugin, applied DADA2^69^ to denoise the data, and trimmed reads in each dataset to the first position with a median quality score under 30. Next, we applied RDP classifier for taxonomic classification according to the GTDB hierarchy^34^. We further filtered samples with less than 2,000 reads or with missing metadata, and similarly removed rare and low abundance taxa, leaving those with abundance >0.05% in at least 5% of the samples. Finally, read counts were normalized to sum to 1 within each sample, resulting in a table of relative abundances.

The obtained FMT data are characterized by triads of samples. Each triad includes one donor sample before FMT, and a pair of recipient samples, one before and one after FMT. Following a previous study that categorized classified taxa into different post-FMT classes^21^ when multiple recipient post-FMT samples were available, they were treated as participating in multiple FMT experiments. Based on the observed composition patterns in donor and recipient samples, taxa were defined as “colonizers” if they appeared in both recipient post-FMT and donor samples but were absent from the recipient pre-FMT sample.

### Calculation of Relative Dispersion Ratio

We calculated the Relative Dispersion Ratio (RDR) between every pair of co-colonizing taxa. The RDR is defined according to the following equation:

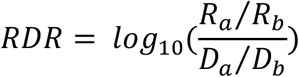

Where, ***R***_*a*_ and ***R***_*b*_ are the relative abundances of the co-colonizing taxa ***a*** and ***b***, respectively, in the post-FMT recipient sample, and ***D***_*a*_ and ***D***_*b*_ are the abundances of these taxa in the donor pre-FMT sample.

### Comparison with in-vitro experiments

We collected results from published co-cultured competition experiments^39^. As the experiments were conducted on several strains from the same genus, we picked a single genus-level representative randomly. We compared the results of the experiments at the end of the 72-hours incubation period to the estimated interactions by RDR. The degree of agreement between the RDR framework and the experimental results was estimated using Cohen’s kappa.

### Comparison with population dynamics

Data from the American Gut Project (AGP) was obtained from the original AGP publication’s accompanying figshare^40^. To allow integration with the FMT studies, we mapped ASV to GTDB genus-level taxonomy^34^. We then removed low-quality samples and filtered rare and low-abundance taxa, following the same criteria as above. We then calculated C-score between every pair of taxa according to the modified formulation of the original work by Stone and Roberts^41,70^, in which the score is standardized by the number of occupied habitats. The following score indicates the degree of segregation between pairs of taxa:

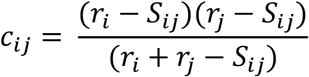

Where, ***c***_*ij*_ is the C-score between taxa ***i*** and ***j***, ***r***_*i*_ and ***r***_*j*_ are the number of samples in which the taxa appear and ***S***_*ij*_ is the number of samples in which the taxa ***i*** and ***j*** appear together.

### Metabolic network analysis

We used the methods described in Levy at el^36^ and in Freilich et al^49^ to investigate seed-sets and metabolic interactions between pairs of taxa. These techniques have been previously utilized to predict metabolic strategies and relationships between bacteria^45^. We employed PICRUST2^47^ to estimate genus-level functional profiles and constructed a representative metabolic network based on the Kyoto Encyclopedia of Genes and Genomes (KEGG)^71^. This network consists of nodes representing compounds and edges representing reactions linking substrates to products, and describe the overall metabolic capacities shared between all members of the genus. From this network, we calculated the seed set, which reflects the nutritional profile of the taxa, as well as the metabolic competition index, which quantified the similarity between the nutritional profiles of the two taxa. To compare the functional capacities of taxa, we used KEGG pathway scores based on their genomic content as calculated using the method described previously^72^. These scores represent the number of KEGG orthology groups (KOs) present in each pathway (with KOs associated with multiple pathways being partitioned between these pathways). We focused only on pathways with at least a two-fold difference (i.e., above 200% or below 50%) between the dominant and the minor pairs. For each such pathway we tested differences in scores between the dominant and the minor partners using Wilcoxon test (FDR p<0.1).

### Machine learning analysis

To predict the composition of the post-FMT community, we constructed a machine learning model for each taxon. We specifically used a random forest regressor that receives as features the pre-FMT abundances of all taxa in the recipient and in the donor. We evaluated the models using 10-fold cross validation and estimated performance using Spearman correlation between the observed and predicted post-FMT abundances. Post-FMT taxa with models that exhibit spearman correlation >0.3 and FDR corrected p-value <0.05 were defined as well-predicted. For these well-predicted taxa, we calculated feature contribution using SHAP values. The machine learning framework was developed using the “Tidymodels” package, random forest models are based on the “Ranger” package^73^, and SHAP calculations were performed by the “FastSHAP” package^74^.

## Supplementary Figures

**Supplementary Figure 1.**
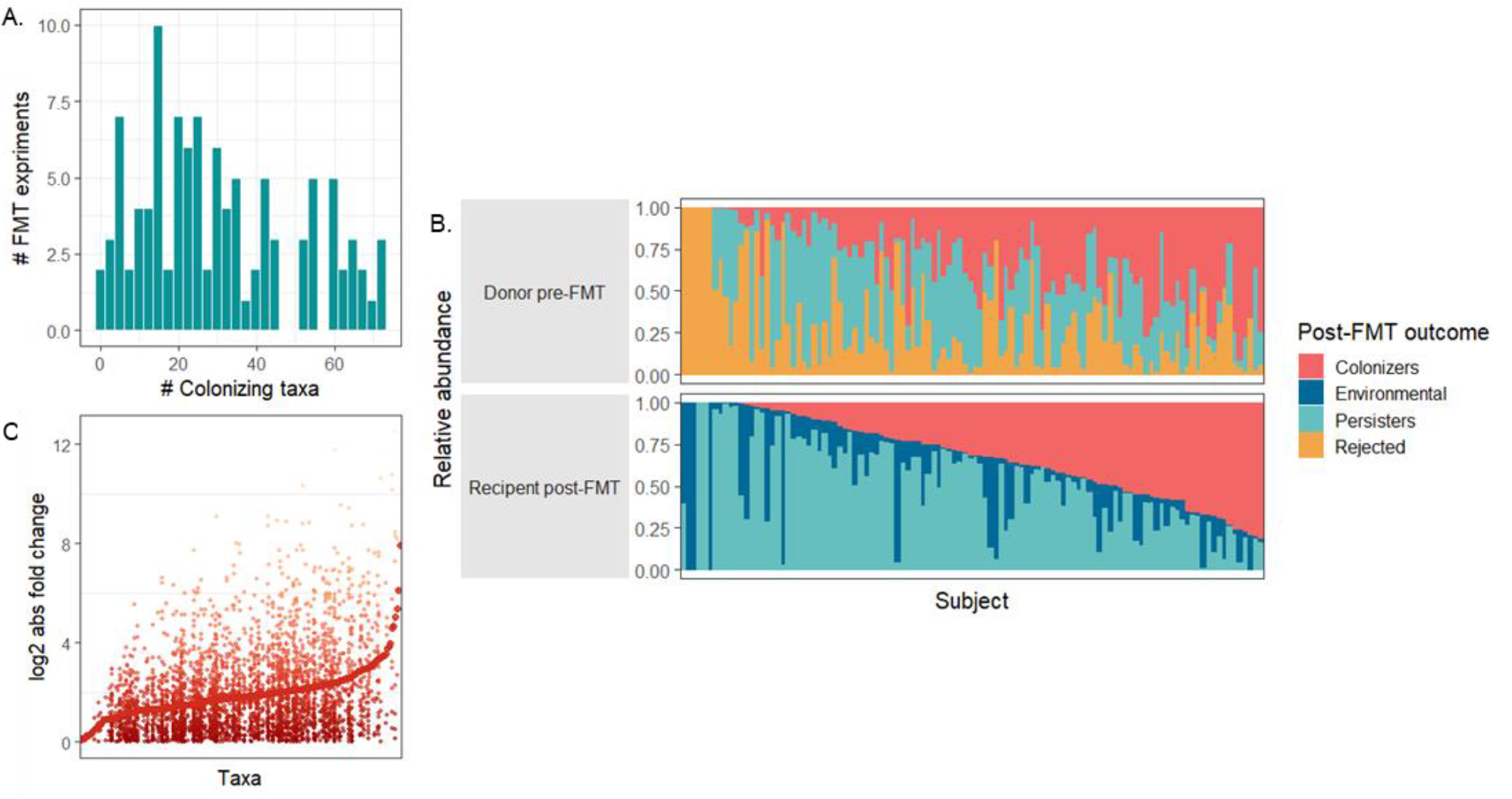
Community composition and colonization frequency following FMT. **A**–Distribution of the number of colonizing taxa observed in each FMT experiment.**B**–Total relative abundance of taxa in the donor pre-FMT and recipient post-FMT samples. Taxa are categorized based on their post-FMT outcomes: colonizing taxa (as described in the main text; highlighted in red), rejected taxa (yellow) that were present in the donor pre-FMT but failed to establish in the recipient post-FMT samples, persistent taxa (light blue) that were identified in both the recipient (pre- and post-FMT) and donor communities, and environmentally acquired taxa (dark blue) that were detected solely in the recipient post-FMT samples. **C**–Colonizing taxa relative abundance fold change (in log2 scale and absolute value) between the donor pre-FMT and the recipient post-FMT environments. Large circles indicate mean fold change for each taxon, and small dots indicate fold change in each FMT experiment.

**Supplementary Figure 2.**
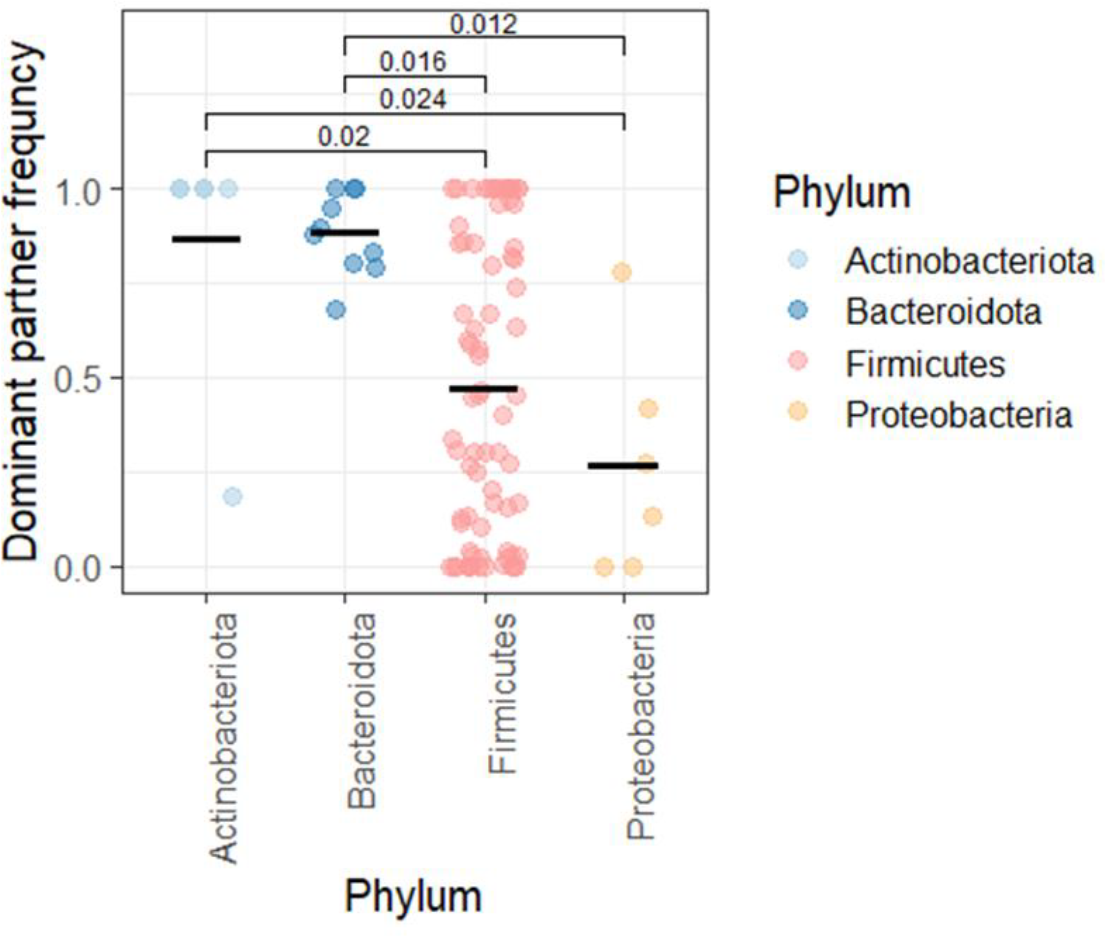
Frequency of dominant partner in various phyla. This figure illustrates the frequency of dominant partnerships based on RDR across different phyla. The y-axis represents the calculated frequency of being the dominant (rather than the minor) partner, while the x-axis displays the various phyla. For clarity and statistical significance, phyla with a sample size of n< 3 were excluded from the figure. To enhance statistical power and consolidate the dataset, we further combined the phyla Firmicutes, Firmicutes_A, and Firmicutes_C. Shown are Wilcoxon test FDR corrected p-value.

## Supplementary Tables

**Supplementary Table 1:** FMT datasets analyzed in our study.

| Study name | Reference | # of patients | # of samples | # of donors | 16s rRNA sequencing region |
| --- | --- | --- | --- | --- | --- |
| autism_kang_2017 | <sup>31</sup> | 18 | 183 | 5 | V4 |
| c_diff_khanna_2017 | <sup>25</sup> | 33 | 123 | 33 | V4 |
| c_diff_seekatz_2014 | <sup>26</sup> | 10 | 30 | 10 | V4 |
| c_diff_zao_2018 | <sup>27</sup> | 11 | 43 | 11 | V3-V4 |
| cancer_baruch_2020 | <sup>32</sup> | 10 | 50 | 2 | V4 |
| crohn_sokol_2020 | <sup>28</sup> | 8 | 56 | 7 | V3-V4 |
| IBD_goyal_2018 | <sup>29</sup> | 21 | 99 | 21 | V4 |
| UC_kump_2017 | <sup>30</sup> | 13 | 40 | 11 | V4 |
| **Total** |  | **124** | **624** | **100** |  |

